# Emergence of population-level feedback control by transposon-plasmid coevolution

**DOI:** 10.1101/2025.10.20.683530

**Authors:** Rohan Maddamsetti, Grayson Hamrick, Yuanchi Ha, Yasa Baig, Jia Lu, Charlotte Lee, Lingchong You

## Abstract

The origins of adaptive functions remain poorly understood, despite considerable interest^1–5^. Here, we report the *de novo* evolution of population-level feedback control in clonal *Escherichia coli* strains containing high-copy plasmids, enabling these strains to express green fluorescent protein (GFP) in response to tetracycline. Selection maintains *tetA*^*+*^ and *tetA*^*–*^ plasmids within single cells, due to negative feedback from the toxic effects of a *tetA-gfp* tetracycline resistance transposon. At high plasmid copy numbers, the intracellular equilibrium of *tetA*^*+*^ and *tetA*^*–*^ plasmids robustly responds to tetracycline through rapid evolutionary dynamics^6–8^. Theory predicts that the GFP response to tetracycline is determined by the covariance between GFP expression and bacterial fitness. Our findings show that small mutational changes can result in large qualitative changes to population-level behavior. Here, the evolution of polymorphic intracellular populations of mobile genetic elements allows host populations to dynamically respond to antibiotic in the environment.

## INTRODUCTION

A central problem for 21^st^ century evolutionary theory is to explain how novel adaptive behaviors emerge in biological systems^9–11^. To address this conceptual issue, it has been proposed that combinations of existing parts may self-assemble and generate qualitatively new structures and functions that natural selection can then modify and refine^3^. According to this hypothesis, a population of simple parts may combine and give rise to unexpected and qualitatively new functions^12^. Experimental evidence of such functional transitions^4,5^ remains limited to a few model systems^13^. In addition, experiments designed to study the evolution of novel functions, such as bet hedging^14^, increased evolvability^15^, and viral host-receptor shifts^16^ are premised on experimental selection for the new trait. The emergence of fractal protein assemblies in the absence of adaptive benefit^17^ suggests that complex protein structures can emerge without selection, but to our knowledge, the emergence of a feedback system from simpler components has not been observed in the laboratory.

Here, we report the discovery of an emergent plasmid-mediated feedback mechanism that allows a clonal bacterial population to respond to tetracycline in the environment. Strikingly, this emergent feedback system can amplify gene expression in response to antibiotics, much like two recent gene circuits that were explicitly designed and constructed by synthetic biologists^18,19^. By contrast with those synthetic systems, however, the intrinsic presence of negative feedback in this evolved system stabilizes intracellular plasmid ratios, without the need for multiple antibiotics or repeated design-build-test cycles for successful function.

## RESULTS

### Emergence of stable coexistence between incompatible plasmids through transposon-plasmid coevolution

We discovered the evolution of stable intracellular coexistence between incompatible plasmids in an experimental system for studying transposon-mediated antibiotic resistance^20,21^. These experiments involved *Escherichia coli* strains harboring a minimal transposon composed of a *tetA* tetracycline resistance gene flanked by short terminal repeats (Figure 1A). The *tetA* mini-transposon is mobilized by an external Tn5 transposase located in the chromosome. In previous work, we showed that the *tetA* transposon rapidly mobilizes from the chromosome onto intracellular plasmids under tetracycline selection^20,21^ (Figure 1B). The *tetA* gene encodes an efflux pump that increases antibiotic resistance, but at the cost of disrupting the membrane potential, causing growth arrest and cell death^22,23^.

**Figure 1:**
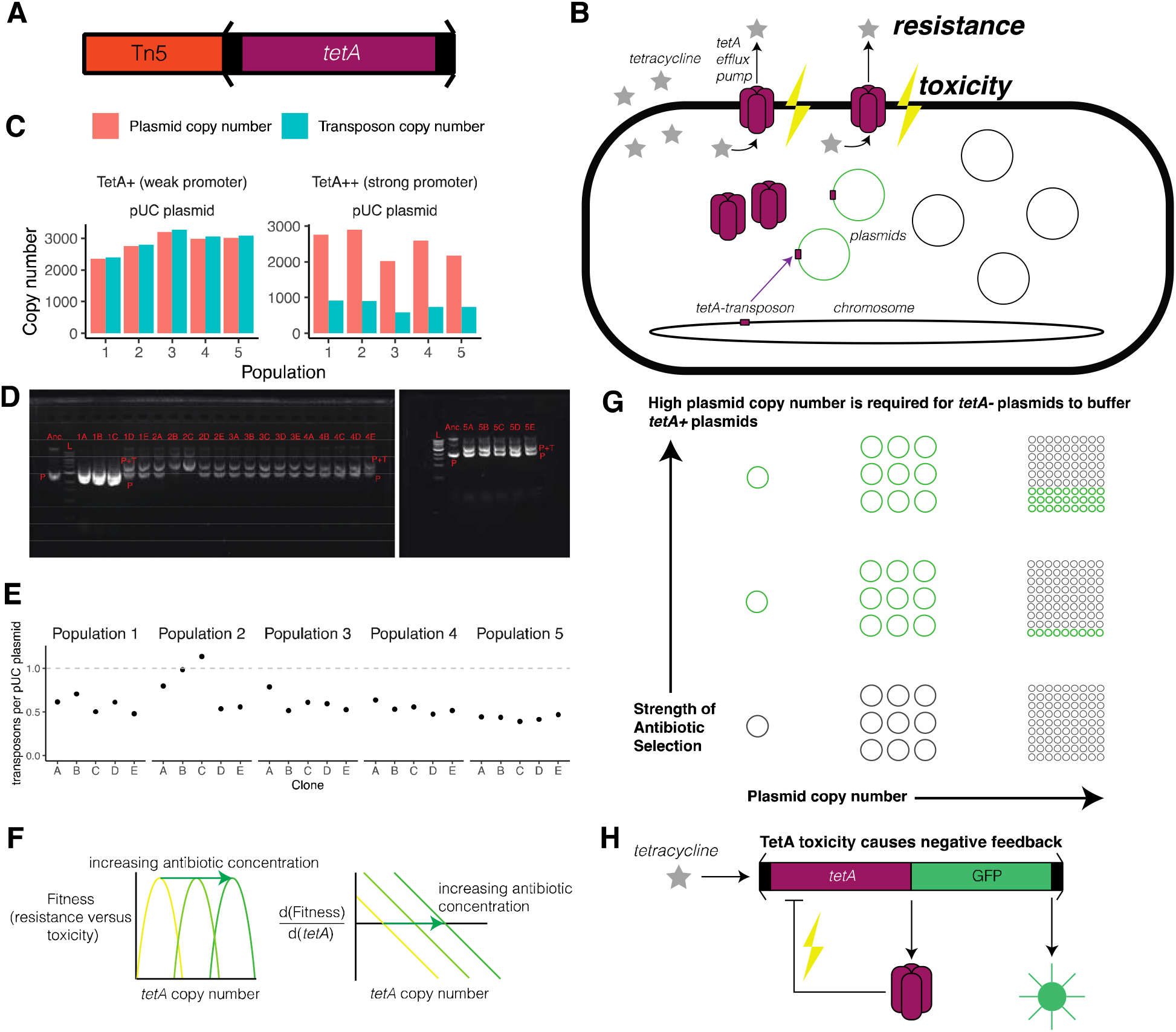
Balancing selection on a toxic *tetA*-transposon drives the *de novo* evolution of plasmid quasispecies in the laboratory. A) Structure of the Tn5 *tetA* mini-transposon. The *tetA* tetracycline resistance gene is flanked by direct repeats. The region within the direct repeats is mobilizable by an external Tn5 transposase. The entire sequence is integrated into the *E. coli* chromosome at the HK022 *attB* site. B) The *tetA* transposon rapidly mobilizes from the chromosome onto intracellular plasmids under tetracycline selection. The TetA efflux pump (in purple) increases antibiotic resistance, but can also result in loss of membrane potential, growth arrest, and cell death (Eckert and Beck 1989). C) *E. coli* populations containing a pUC plasmid were evolved for nine days under increasing tetracycline selection, starting from clonal ancestral strains containing a Tn5 *tetA* mini-transposon containing either a weak promoter controlling *tetA* expression (TetA+) or a strong promoter (TetA++). *tetA* and pUC plasmid copy numbers in the endpoint population samples were measured by genome sequencing. D) DNA electrophoresis demonstrates coexistence of *tetA*^*–*^ and *tetA*^*+*^ pUC plasmid variants within single clones from the endpoint populations of the evolution experiment. Five random clones (A through E) were picked from each population (1 through 5), and purified plasmids were run on 1% agarose gels. “Anc.” denotes the ancestral strain before evolution, “L” denotes the DNA marker ladder, “P” denotes the plasmid (without transposon), and “P+T” denotes the plasmid with transposon. E) qPCR demonstrates coexistence of *tetA*^*–*^ and *tetA*^*+*^ pUC plasmid variants within single clones from the endpoint populations of the evolution experiment, confirming that plasmids were not saturated with transposons. F) The optimal number of *tetA* copies carried on intracellular plasmids is tunable by antibiotic selection, due to the balance between resistance and toxicity caused by *tetA*. Balancing selection leads to a stable intracellular equilibrium between *tetA*^*–*^ and *tetA*^*+*^ plasmid variants. G) High plasmid copy number is required to maintain an intracellular balance between *tetA*+ and *tetA*– plasmids. At low and medium plasmid copy numbers, stochastic partitioning of *tetA*+ and *tetA*– plasmids eventually leads to daughter cells that either contain only *tetA*+ or only *tetA*– plasmids, the latter going extinct under strong antibiotic selection. H) Given the emergence of population-level negative feedback control due to transposon-plasmid coevolution, the expression of a gene of interest (such as green fluorescent protein) linked to *tetA* can be controlled by modulating environmental tetracycline concentrations.

We carried out 9-day evolution experiments with *E. coli* DH5α and sequenced endpoint populations resistant to 50 μg/mL tetracycline, varying replication origin (and thus plasmid copy number), the presence of active transposase, and the basal expression of the *tetA* resistance gene^21^. When *tetA* expression was driven by a strong promoter (TetA++), final *tetA-*transposon copy numbers were much lower than pUC plasmid copy numbers; by contrast, when *tetA* expression was driven by a weak promoter (TetA+), the *tetA*-transposon saturated high copy number pUC plasmids across all experimental populations (Figure 1C). In treatments containing a medium copy number p15A plasmid, the TetA++ *tetA*-transposons saturated the p15A plasmid (Supplementary Figure S1). In addition, *tetA*-transposon copy numbers never changed in a no-tetracycline control treatment, indicating that the combination of tetracycline selection, high copy number pUC plasmids, and strong TetA++ expression was required for *tetA-*transposon copy numbers < high pUC copy numbers (Supplementary Figure S1).

We hypothesized that the toxicity of TetA++ expression^22^ prevented the *tetA-* transposons from saturating high copy number pUC plasmids, leading to coexistence of *tetA*^*+*^ and *tetA*^*–*^ plasmids within single cells (Hypothesis 1). Since these data represent measurements across the bacterial population, these data could alternatively be explained by the co-existence of different bacterial subpopulations containing either *tetA*^*–*^ or *tetA*^*+*^ pUC plasmids (Hypothesis 2). To distinguish between these hypotheses, we isolated five clones (labeled A through E) from each of the five replicate TetA++ populations, purified pUC plasmids from these clones, and ran the purified plasmids on 1% agarose gels (Figure 1D). As a control, we ran samples of the pUC plasmid from the ancestral strain of the TetA++ treatment prior to the evolution experiment. Under Hypothesis 1, each clone should contain *tetA*^*–*^ and *tetA*^*+*^ pUC plasmids, resulting in two bands per clone. By contrast, if Hypothesis 2 were correct, then each clone should contain either a *tetA*^*–*^ and *tetA*^*+*^ pUC plasmid, resulting in one band per clone.

The left gel in Figure 1D shows the ancestral pUC plasmid, the molecular ladder, and the evolved pUC plasmids purified from the five clones from the first four evolved replicates of the TetA++ pUC treatment. The right gel in Figure 1D shows the molecular ladder, the ancestral pUC plasmid, and the evolved pUC plasmids purified from the five clones from the fifth evolved replicate of the TetA++ pUC treatment. Clones A, B, and C from the first replicate show bright bands on the gel. Clones D and E show two bands. The lower molecular weight band corresponds to *tetA*^*–*^ pUC plasmids without the transposon, and the higher molecular weight band corresponds to *tetA*^*+*^ pUC plasmids containing the transposon. The next lane corresponds to clone A of the second TetA++ pUC replicate, again showing two distinct bands. Clones B and C from the second replicate show the higher molecular weight band, but not the smaller band. All clones from replicate populations 3 and 4 on the left gel show two distinct bands, and all clones from replicate population 5 in the right gel show two distinct bands. These data indicate the presence of multiple pUC variants in clones isolated from all five TetA++ pUC replicate populations, which supports Hypothesis 1: *tetA*^*–*^ and *tetA*^*+*^ pUC plasmid variants coexist within TetA++ pUC clones. As a validation, qPCR was used to measure the ratio of transposon copies to plasmid copies in each of these clones. 23 out of 25 clones had *tetA* copy numbers lower than pUC plasmid copy numbers, indicating the evolution of stable plasmid quasispecies consisting of coexisting *tetA*^*–*^ and *tetA*^*+*^ pUC plasmid variants within single clones (Figure 1E).

Based on these experimental findings, we hypothesized that the balance between *tetA*^*–*^ and *tetA*^*+*^ pUC plasmid variants within single cells could be tuned by modulating tetracycline concentrations. According to this conceptual model, the optimal *tetA* copy number is a function of the environmental tetracycline concentration. When *tetA* copy numbers are too low, the bacteria have insufficient resistance. When *tetA* copy numbers are too high, TetA protein toxicity leads to cellular growth arrest and death. The balance of these effects leads to a stable intracellular equilibrium between *tetA*^*–*^ and *tetA*^*+*^ plasmid variants that can be modulated by tetracycline selection (Figure 1F).

The existence of a stable intracellular equilibrium depends on the plasmid copy number, or in other words, the size of the intracellular plasmid population. Sufficiently high plasmid copy numbers are required to maintain an intracellular balance between *tetA*^*+*^ and *tetA*^*–*^ plasmids. Given low or medium plasmid copy numbers, stochastic partitioning (genetic drift) eventually leads to daughter cells that either contain only *tetA*^*+*^ or only *tetA*^*–*^ plasmids. In this scenario, strong antibiotic selection favors cells that only contain *tetA*^*+*^ plasmids, all else being equal (Figure 1G).

A key prediction of the conceptual model shown in Figure 1F and Figure 1G, is that the intracellular balance of *tetA*^*–*^ and *tetA*^*+*^ plasmid variants can track environmental tetracycline concentrations, once plasmid copy numbers cross a critical threshold. We therefore hypothesized that population-level negative feedback control emerged in our evolution experiments, without explicit design or construction. By linking a gene of interest, such as green fluorescent protein (*gfp*), to *tetA* copy number, it should be possible to control population-level gene expression through the negative feedback imposed by *tetA* toxicity (Figure 1H). In other words, negative frequency-dependent selection within the level of the intracellular plasmid population should allow single bacterial clones to rapidly change population-level gene expression in response to tetracycline.

### Theory demonstrates conditions for the emergence of population-level negative feedback control

#### Mathematical model

We built a mathematical model to examine the conditions in which population-level negative feedback control would emerge (Figure 2). To simplify the mathematical analysis, we assume that the total number of plasmids per cell are fixed. We also assume that plasmid segregation rates are much higher than genomic mutation rates, such that the effects of *de novo* mutations in the chromosome or plasmid can be largely ignored. Single-nucleotide mutation rates and small-indel mutation rates in wildtype *E. coli* are on the order of 10^−10^ per site per generation, and 10^−11^ per site per generation, respectively^24^. Scaling by the length of the *E. coli* genome (~5×10^6^ nt) yields a genomic mutation rate of 10^−4^ per genome per generation. Transposition rates in wildtype *E. coli* are on the order of 10^−5^ per transposon per genome per generation^25,26^. By contrast, binomial sampling of *tetA*^*+*^ and *tetA*^−^ plasmids from a parent cell containing a 50:50 ratio of *tetA*^*+*^ to *tetA*^−^ plasmids has a probability of *tetA*^+^ copy number change of 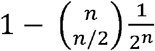 per generation, where *n* is the plasmid copy number. When *n* = 2, the probability of *tetA*^*+*^ copy number change by binomial sampling is 0.5 per generation; when *n* = 100, the probability of *tetA*^*+*^ copy number change per generation by binomial sampling is 0.92 per generation, which exceeds the genomic mutation rate by several orders of magnitude. This back-of-the-envelope calculation is conservative, because a binomial model of random plasmid segregation underestimates the heterogeneity seen in populations of cells containing high-copy-number plasmids^27^.

**Figuree 2.**
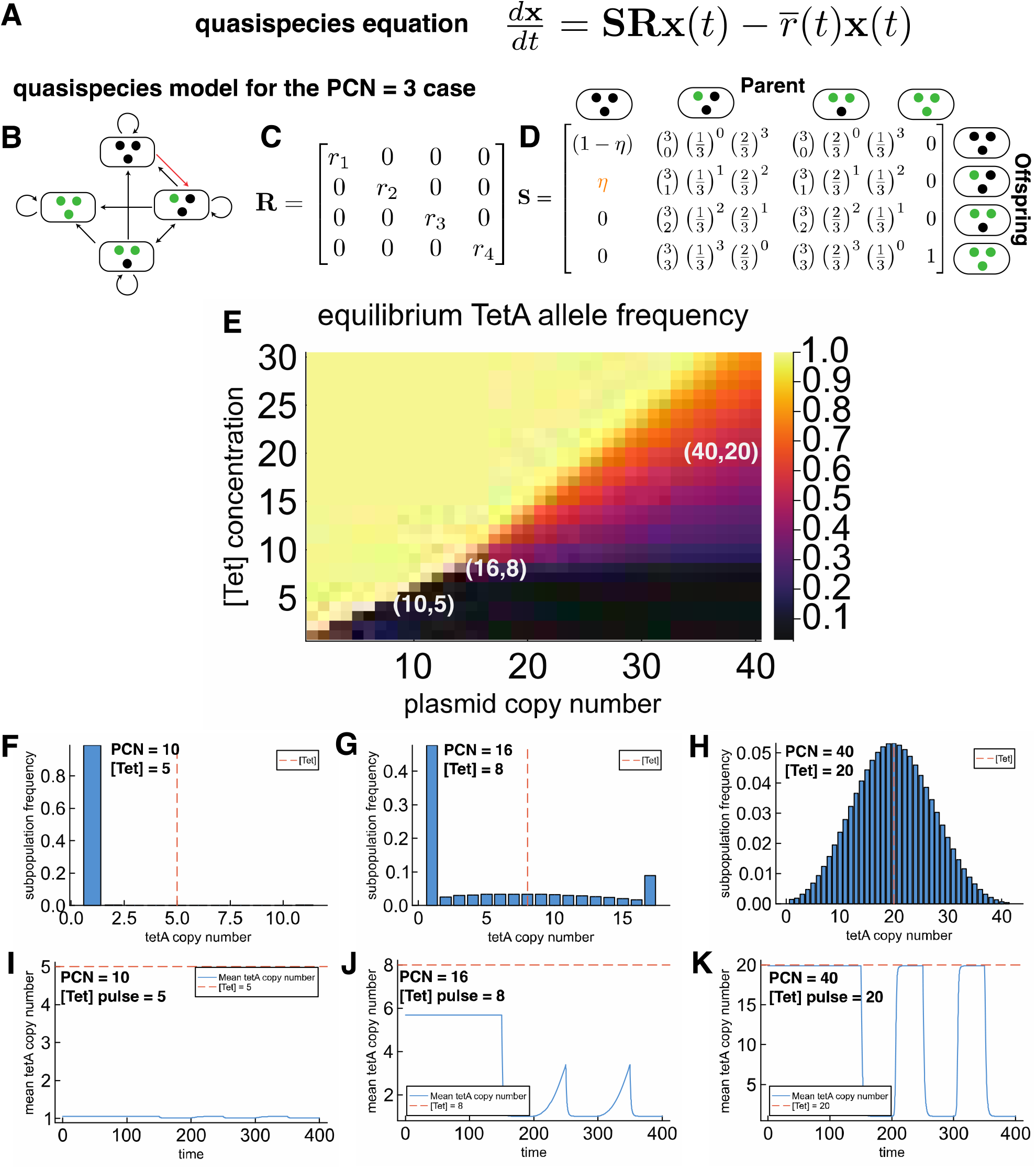
Theory and simulations demonstrate how population-level feedback control can emerge in bacteria. A) Matrix algebra form of the quasispecies equation. B) Diagram of the quasispecies model in the PCN = 3 case. The orange arrow indicates the transposition of the *tetA*-transposon from the chromosome to a single plasmid. C) R matrix representing fitnesses (growth rates) for each cellular state for the PCN = 3 case. D) S matrix representing plasmid segregation and transposition dynamics for the PCN = 3 case. The orange entry represents the rate at which the *tetA*-transposon jumps from chromosome to plasmid. E) Phase diagram shows the parameter space of tetracycline concentrations [Tet] and plasmid copy numbers required for stable intracellular coexistence of *tetA*^*+*^/*tetA*^−^ plasmids in a constant environment. The stability of the intracellular *tetA*^*+*^ allele frequency equilibrium increases with plasmid copy number. The yellow region corresponding to intracellular allele frequency = 1, meaning that only *tetA*^*+*^ plasmids are present. The black region corresponding to intracellular allele frequency = 0, meaning that only *tetA*^−^ plasmids are present. The colorful region corresponds to intermediate intracellular allele frequencies at which *tetA*^*+*^ and *tetA*^−^ plasmids have a stable intracellular coexistence. F) Stationary distribution of *tetA* copy numbers when PCN = 10 and [Tet] = 5. G) Stationary distribution of *tetA* copy numbers when PCN = 16 and [Tet] = 8. H) Stationary distribution of *tetA* copy numbers when PCN = 40 and [Tet] = 20. I) Population dynamics in response to pulses of [Tet] = 5 when PCN = 10. J) Population dynamics in response to pulses of [Tet] = 8 when PCN = 16. K) Population dynamics in response to pulses of [Tet] = 20 when PCN = 40. Tunable population dynamics emerge in the *tetA*^*+*^/*tetA*^−^ coexistence phase.

We model the bacterial population as a vector **x**(*t*) indexed by *tetA* copy numbers, ranging from 1 (the single copy found in the chromosome) to *n+1*, where *n* is the plasmid copy number (PCN). The population dynamics are determined by the quasispecies equation^1,2,28,29^ shown in Figure 2A. The basic structure of the model is shown in Figure 2B, using the PCN = 3 case as an example. The growth rates (fitness) for each *tetA* copy number class are defined in the matrix **R** (Figure 2C). The growth rates in the **R** matrix depend on *tetA* copy number and [Tet], the environmental concentration of tetracycline (Methods). For simplicity and ease of interpretation, we parameterize the growth rates as a Gaussian function, such that the optimal *tetA* copy number (TCN) class is equal to [Tet], after multiplication by the appropriate scale factor to change units. Decreasing [Tet] has the effect of translating the Gaussian fitness function to the left, while increasing [Tet] translates the Gaussian fitness function to the right. The total number of offspring (i.e., total growth) per unit time is determined by the matrix-vector product **Rx**(*t*). The offspring per parent *tetA* copy number class are partitioned into daughter *tetA* copy number classes based on binomial sampling, as defined by the **S** matrix (Figure 2D). In addition, we assume that the lowest *tetA* copy number class (one copy on chromosome) generates mutants with an additional *tetA* copy on a plasmid at a transposition rate η. The relevant entry **S**_21_ showing the transition from 1 *tetA* copy to 2 *tetA* copies is shown in orange in Figure 2D. The composition of growth and transposition η*r*_*1*_ is shown in the model diagram in Figure 3B as an orange arrow. The composition of the **S** and **R** matrices describes mutation-selection dynamics: the **R** matrix represents differential growth among *tetA* copy number classes (i.e. selection), and the **S** matrix represent plasmid segregation and *tetA-*transposon transposition (“mutation” in the generalized sense of transitions between *tetA* copy number classes due to plasmid segregation).The whole population grows at the mean growth rate (i.e. mean fitness) 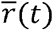. The final term of quasispecies model represents dilution of the population at rate 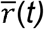 to balance out overall population growth, to maintain a constant population density over time.

**Figuree 3.**
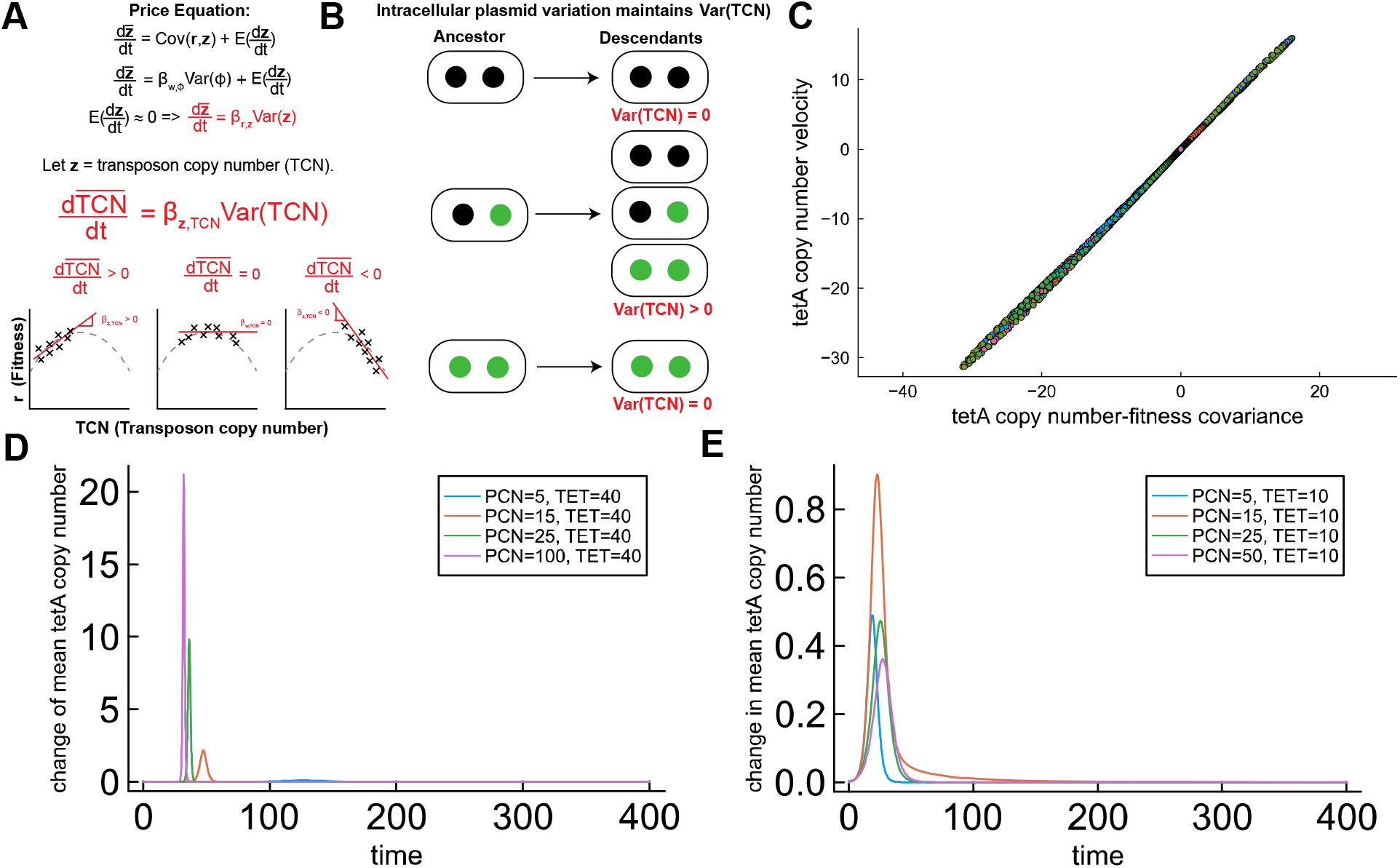
The response of the feedback system is determined by covariance between *tetA* copy number and bacterial fitness. A) The Price equation implies that instantaneous change in TCN depends on the slope of the linear regression of fitness on TCN in plasmid quasispecies and the variance in TCN in the quasispecies. B) Balancing selection maintains population variance in TCN. C) 100 random pairs of tetracycline concentration and PCN parameters show that the rate of mean *tetA* copy number is equal to the covariance between bacterial fitness and TCN in the mathematical model, as predicted by the Price equation. D) Under strong selection ([Tet] = 40), the covariance between fitness and *tetA* copy number is maximized at high PCN (PCN = 100); in this case, system response is maximized at high PCN. E) Under weaker selection ([Tet] = 10) the covariance between fitness and *tetA* copy number is maximized at medium PCN (PCN = 15); in this case, system response is maximized at medium PCN and decreases at high PCN.

#### Properties of the quasispecies model

The quasispecies equation has elegant properties that shed insight into the key principles of our experimental system. First, it has an intuitive physical interpretation. In this context, it models a bacterial population growing exponentially, with a large excess of nutrients, in an idealized turbidostat^30^ that instantaneously increases its outflow rate as the population increases in mean fitness, thus maintaining a constant population density. Second, it is easy to diagonalize the matrix product **SR** numerically and calculate its eigenvectors and associated eigenvalues. The right eigenvector associated with the dominant eigenvalue, when normalized, is equal to the stationary distribution of the quasispecies, assuming fixed [Tet] and thus a fixed **R** matrix. The dominant eigenvalue is equal to the mean population fitness of this stationary quasispecies^1,2,28^. These facts let us immediately calculate the stationary distribution of the quasispecies at a fixed [Tet] concentration across a wide range of [Tet] and PCN values. This allows us to easily calculate a phase diagram that describes how the stability of the intracellular coexistence between *tetA*^*+*^ and *tetA*^−^ plasmids and the shape of the stationary distribution change with [Tet] and PCN as control parameters (Figure 2E).

#### Phase transitions in the plasmid quasispecies model

The phase diagram in Figure 2E shows three distinct phases in the parameter space set by PCN and [Tet]: first, a state in which the *tetA*^*+*^ plasmids are monomorphic at the stationary state (the yellow region corresponding to intracellular plasmid allele frequency = 1 in the phase diagram); second, a state in which *tetA*^−^ plasmids are monomorphic at the stationary state (the black region corresponding to intracellular plasmid allele frequency = 0 in the phase diagram); and third, an intermediate phase in which *tetA*^*+*^ and *tetA*^−^ plasmids have a stable intracellular coexistence (the colorful region corresponding to intermediate intracellular plasmid allele frequencies). Two kinds of phase transitions can lead to the intracellular coexistence of *tetA*^*+*^ and *tetA*^−^ plasmids. First, when plasmid copy numbers are high, increasing [Tet] leads to the *tetA*^*+*^/*tetA*^−^ coexistence phase. Second, when [Tet] is high, increasing PCN leads to the *tetA*^*+*^/*tetA*^−^ coexistence phase.

The coexistence phase is demarcated by positive Shannon entropy (Supplementary Figure S2A), *tetA* copy number variance (Supplementary Figure S2B), and fitness variance (Supplementary Figure S2C). This coexistence phase shows the region of PCN and [Tet] parameter space in which *tetA*^*+*^/*tetA*^−^ plasmids have a stable intracellular allele frequency, based on intracellular “negative-frequency dependent selection”, based on the trait encoded in the intracellular population (tetracycline resistance, in this case).

An example of a quasispecies outside of the coexistence phase is shown in Figure 2F, which shows the stationary population distribution when PCN = 10 and [Tet] = 5. The population is almost entirely comprised of cells that only contain *tetA*^−^ plasmids. Figure 2G shows a quasispecies just across the phase transition threshold, at PCN = 16 and [Tet] = 8. Figure 2H shows a quasispecies well within the coexistence phase, at PCN = 40 and [PCN] = 20. This last case illustrate how high PCN allows a single bacterial cell to generate daughter cells containing diverse configurations of intracellular *tetA*^*+*^ and *tetA*^−^ plasmids, generating population-level diversity as a stationary state.

When plasmid copy numbers are sufficiently high, population-level gene expression dynamics can be controlled by modulating tetracycline concentrations and switching the population in and out of the *tetA*^*+*^/*tetA*^−^ coexistence phase. When PCN = 10 and the population experiences alternating pulses of [Tet] = 5 and [Tet] = 0, the population barely responds (Figure 3I). When PCN = 16 and the population experiences alternating pulses of [Tet] = 8 and [Tet] = 0, the population is able to respond, but not fast enough for the mean to each the optimal PCN = 8 state indicated by the dashed red line (Figure 3J). When PCN = 40 and the population experiences alternating pulses of [Tet] = 20 and [Tet] = 0, the population rapidly switches back and forth and is easily able to reach the optimal PCN = 20 state (Figure 2K).

This mathematical analysis demonstrates four key points of our proposed mechanism (Figure 1FGH). First, balancing selection maintains genetically diverse intracellular plasmid populations within single cells, due to the toxic effects of a *tetA*-*gfp* tetracycline resistance transposon. Second, the intracellular balance of *tetA*^*+*^ and *tetA*^−^ plasmids responds to tetracycline selection, provided that plasmid copy number is sufficiently high. Third, the stability of the balancing selection equilibrium and the magnitude of the gene-expression response increases with plasmid copy number. Fourth, when single bacterial clones are isolated from the coexistence phase, the distribution of plasmids within cells serves as an internal model of the environment, allowing the entire bacterial population to operate as a feedback control system that can track environmental antibiotic concentrations.

#### The tetA^+^/tetA^−^ coexistence phase is suboptimal in a constant environment

Our mathematical analysis reveals another fact: the *tetA*^*+*^/*tetA*^−^ coexistence phase is always suboptimal in a constant environment. Recall that the optimal fitness is reached in our model when *tetA* copy number (TCN) = [Tet]. This means that fitness is maximized when the entire population has TCN = [Tet]. By comparing the ratio of mean population fitness to maximum population fitness (i.e., fitness when TCN = [Tet]), our model shows that plasmid heterogeneity reduces mean population fitness by generating suboptimal daughter cells with TCN ≠ [Tet] (Supplementary Figure S3). In a constant environment, reducing population variance around the optimum maximizes fitness.

### The Price equation determines response properties of the plasmid quasispecies

Biological engineers have built coupled systems of intracellular plasmids to control and amplify gene expression in bacterial populations^18,19^. However, fundamental properties of such systems have not been described, due to limited communication between evolutionary and synthetic biologists. Our theoretical framework shows how a fundamental law called the Price equation has implications for engineering gene expression with intracellular plasmid populations.

The Price equation is the fundamental theorem of evolution^31^. Straightforward proofs of both discrete-time and continuous-time theorems are given by Liorsdottir and Pachter^32^; we use the continuous-time theorem here. Briefly, the Price equation states that the instantaneous rate of change of the mean of any trait depends on the covariance between that trait and fitness, and the expected change of trait values over time. Let **z** denote a fixed vector of trait values. In this case, **z** represents *tetA* copy number classes ranging from 1 (the single copy found in the chromosome) to *n+1*, where *n* is the plasmid copy number (PCN). Following Liorsdottir and Pachter^32^, the continuous Price equation is

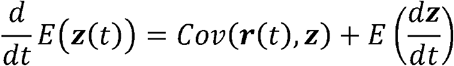

where **r**(*t*) is a vector of growth rates (or fitnesses) of the trait value classes, and *E* and *Cov* denote expected value and covariance, respectively. **r**(*t*) is a function of time because we assume that the growth rates depend on antibiotic concentrations that may vary over time. While the vector **z** is fixed, we use the notation 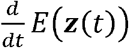 to indicate that its expected value changes with time: *E*(*z* (*t*)) = ∑_*i*_ *z*_*i*_ *x*_*i*_ (*t*) where the vector x (*t*) denotes the frequencies of cells with different *tetA* copy numbers, per the quasispecies equation (Figure 3). Page and Nowak (2002) show that combining the quasispecies equation with the continuous-time Price equation results in the following expanded Price equation^29^:

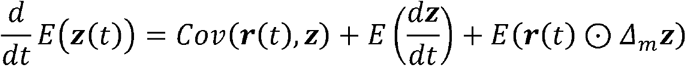

Here, the final term *E* ( *r* (*t*) ⊙ Δ_*m*_ *z* describes transitions among types (in our model, due to plasmid segregation during cell division), with Δ_m_*z*=∑_*j*_ *S*_*ji*_ (*z*_*j*_ - *z*_i_) denoting expected change in trait value when mutating from *tetA* copy number class *i*. ⊙ The symbol represents the element-wise Hadamard product between vectors of the same dimension.

In our model, 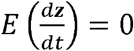 because the trait vector **z** is defined by *tetA* copy classes (the trait frequency vector **x**(*t*) changes over time, but the trait values **z** are fixed)^32^. It follows that:

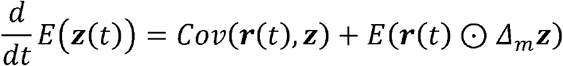

Suppose that *E* (*r* (*t*) ⊙ Δ_*m*_ *z*) ≈ 0. This means that offspring do not systematically deviate from their parents in fitness, which implies no transmission bias in the plasmids. This condition holds in our model because the binomial transitions in the **S** matrix are symmetric, leading to an expected change of zero. The only asymmetric mutational pressure is the rate at which transposons jump from chromosomes to plasmid (the entry **S**_21_), which we assume to be small. Because **z** is the vector of *tetA*-transposon copy number classes *TCN*, we deduce that:

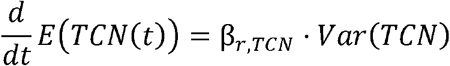

because the regression coefficient β_*r,z*_ is defined as 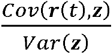. This result gives the instantaneous change of mean TCN in the population and shows that this rate depends on the slope of the linear regression of fitness on TCN in the cloud of genotypes determined by the plasmid quasispecies, and the variance in TCN in the quasispecies. This result has a simple geometric interpretation^33^ (Figure 3A). Under strong directional antibiotic selection, increasing plasmid copy number increases variance in TCN, and accelerates the response of the population (Figure 3A). Balancing selection maintains intracellular plasmid variation and therefore maintains variance in TCN (Figure 3B).

We numerically verified the Price equation in our model by choosing 1000 different random sets of PCN and [Tet] parameters; in all cases, the velocity at which *tetA* copy number changes in each population corresponds one-to-one with the covariance between *tetA* copy number and fitness (Figure 3C). We also checked the correspondence for the specific parameters sets examined in the phase diagram shown in Figure 3. These results are shown in Supplementary Figure S4. In all cases, the Price equation exactly determines change in mean *tetA* copy number. When only the covariance term is considered, the correspondence is almost exact: a small error in seen in the case that PCN = 10 and [Tet] = 5, due to the small effect of the asymmetric mutation term corresponding to transpositions from chromosome to plasmid. As PCN increases, this error becomes negligible, and the covariance between *tetA* copy number and fitness is sufficient to describe the dynamics of *tetA* copy number change (Supplementary Figure S4).

The linear regression interpretation of the Price equation implies that increasing PCN increases the velocity of the feedback system response to changes in tetracycline, as long as the slope of the linear regression does not decrease with higher PCN. In our model, when [Tet] = 40, increasing PCN speeds the response (Figure 3D). At lower [Tet] concentrations, however, the slope of the linear regression can *decrease* for populations with higher PCN, compared to populations with lower PCN that are better able to reach the optimal TCN. One such example is shown in Figure 3E, where plasmid quasispecies with PCN = 50 have a slower response to [Tet] = 10 selection than a plasmid quasispecies with PCN = 15.

These analytical results show engineering implications of the Price equation. The Price equation dictates that the dynamics of an arbitrary engineered trait carried on intracellular plasmid populations depends on the covariance between that trait and bacterial fitness.

### Emergence of clonal bacterial strains that express population-level pulses of GFP in response to pulses of tetracycline

Our mathematical modeling illustrates our hypotheses that 1) stable coexistence between incompatible plasmids emerges by transposon-plasmid evolution in our experiments, when PCN is sufficiently high, and 2) single clones with sufficiently high PCN would be able to change population-wide gene expression in response to changing environmental tetracycline concentrations.

To test these hypotheses, we cloned GFP as a transcriptional reporter downstream of *tetA* in the mini-Tn5 transposon (Figure 4A), and then integrated the construct into the *E. coli* chromosome as before (Methods). Replicate cultures were passaged for one day in LB + 5 μg/ml tetracycline, with high-copy pUC plasmids, high-copy CloDF13 plasmids, medium-high ColE1 plasmids, medium-copy p15A plasmids, or low-copy pSC101 plasmids. Cultures were diluted and plated on selective agar to isolate single clones.

**Figuree 4.**
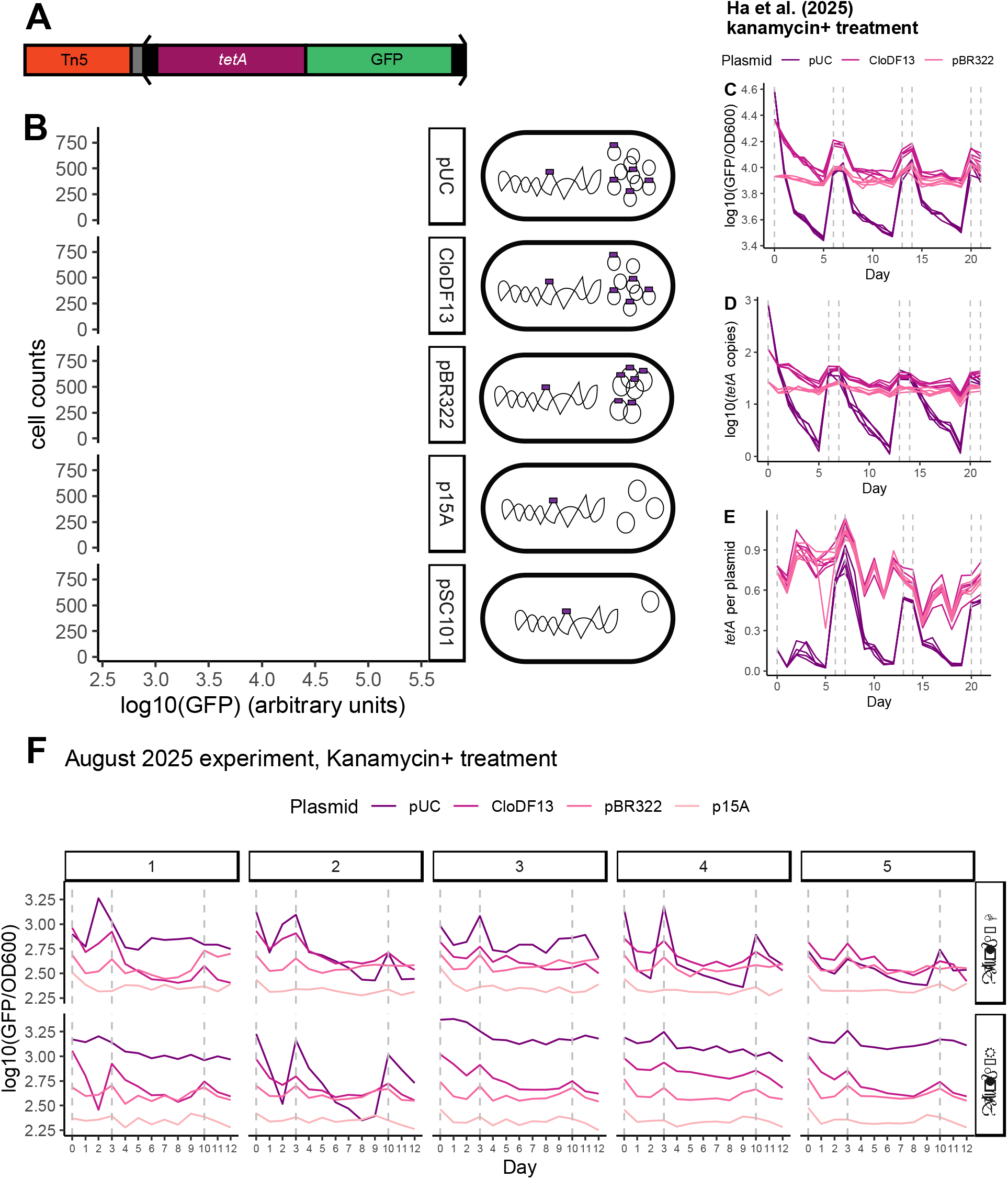
Emergence of clonal bacterial strains that express population-level pulses of GFP in response to pulses of tetracycline. A) Structure of the miniTn5-TetA-GFP transposon. The *tetA* tetracycline resistance gene, followed by a GFP transcriptional reporter is flanked by direct repeats. The region within the direct repeats is mobilizable by an external Tn5 transposase. A random barcode sequence is located outside of the transposable region as well. The entire sequence is integrated into the *E. coli* chromosome at the HK022 *attB* site. B) Flow cytometry shows the amplification and variability of GFP distribution in bacterial clones containing the *tetA-gfp*-transposon and a high copy-number pUC plasmid, a high copy-number CloDF13 plasmid, a medium-high copy-number ColE1 plasmid, a medium copy-number p15A plasmids or a low copy-number pSC101 plasmid. C) Experiments carried out by Ha et al. (2025) show that GFP expression (normalized by cell density) rapidly evolves in response to changes in tetracycline concentration, but only when plasmid copy numbers are sufficiently high. Dashed gray lines indicate timing of tetracycline pulses. D) Experiments carried out by Ha et al. (2025) show that qPCR shows that that the intracellular *tetA* copy on plasmids tracks environmental tetracycline concentrations when plasmid copy numbers are sufficiently high. Dashed gray lines indicate timing of tetracycline pulses. E) Experiments carried out by Ha et al. (2025) show that qPCR shows that that the intracellular *tetA* allele frequency on plasmids tracks environmental tetracycline concentrations when plasmid copy numbers are sufficiently high. Dashed gray lines indicate timing of tetracycline pulses. F) Follow-up validation experiment on clones isolated through a second bottleneck shows emergence of clonal bacterial strains that express population-level pulses of GFP in response to pulses of tetracycline. Dashed gray lines indicate timing of tetracycline pulses.

By picking single clones for our evolution experiments, we can rule out the alternative hypothesis that pulses in GFP expression in response to pulses of tetracycline were caused by competing subpopulations of cells containing monomorphic *tetA*^*+*^ or *tetA*^−^ plasmids; instead, any *tetA*^*+*^/*tetA*^−^ variation must be caused by coexisting *tetA*^*+*^/*tetA*^−^ plasmids within the clone picked for downstream experiments.

DNA electrophoresis showed that tetracycline selection resulted in clones with coexisting *tetA*^*+*^/*tetA*^−^ plasmids in the case of the pUC and CloDF13 plasmids (Supplementary Figure S5). The gel data for CloDF13 indicated the presence of *tetA*^*+*^ plasmids. The p15A clone only contained a *tetA*^−^ plasmid, and the concentration of the pSC101 plasmid was too low to see on agarose gels (Supplementary Figure S5). Flow cytometry showed amplifications of GFP expression and increased GFP variance with increasing plasmid copy number in these clones (Figure 4B).

#### Experiments reported by Ha et al. (2025) demonstrate that high-copy plasmids are necessary for tunable transposon dynamics

Under our hypothesis, we should see only tunable dynamics in clones containing with sufficiently high copy number plasmids to maintain a balance between *tetA*^*+*^/*tetA*^−^ plasmids. The experiments reported in a companion paper^34^ support this prediction. Specifically, Ha et al. (2025) conducted evolution experiments with these pUC, CloDF13 and pBR322 (ColE1) plasmid clones, in which a pulse of 5 μg/ml tetracycline was used for two days of overnight growth, followed by several days of growth without tetracycline. These cycles were carried out by Ha et al. 3 times (Figure 4C, 4D and 4E).

In an experimental treatment in which kanamycin selection was used to maintain the plasmids over time, the pUC clone showed large increases in GFP expression in response to tetracycline selection, with a loss of GFP expression after days of growth in tetracycline-free media (Figure 4C). The change of GFP expression is paralleled by changes in the *tetA-gfp* transposon copy number (Figure 4D). Importantly, the ratio of *tetA* copy number to plasmid copy number (i.e., the intracellular frequency of *tetA*^*+*^ plasmids) showed a large response to changes in tetracycline concentration for the pUC clone but not for the CloDF13 and pBR322 clones (Figure 4E). In an experimental treatment without kanamycin selection, the pUC clone, the CloDF13 clone, and the pBR322 clone all show large changes in GFP expression (Supplementary Figure S6A) and *tetA-gfp* copy number (Supplementary Figure S6B) in response to changes in tetracycline selection. The frequency of *tetA*^*+*^ plasmids responds to tetracycline selection in the populations founded by the pUC clone, but not the CloDF13 and pBR322 clones (Supplementary Figure S6C).

Together, the data reported by Ha *et al*. demonstrate 1) the emergence of tunable population-level gene-expression dynamics in intracellular plasmid quasispecies, once plasmid copy numbers are sufficiently high, and 2) the balance of *tetA*^*+*^*/tetA*^−^ plasmids can be modulated by tetracycline selection when plasmid copy numbers are sufficiently high, as seen for the pUC clone but not the CloDF13 or pBR322 clones.

While our mathematical model assumes that PCN are fixed, these data indicate that PCN can also respond to tetracycline selection. The consequences of variable PCN are explored in the paper by Ha et al.^34^ Regardless, these data show that the equilibrium between *tetA*^*+*^*/tetA*^−^ plasmids is tunable when PCN becomes sufficiently high (Figure 4D and Supplementary Figure S6B), as predicted by our mathematical model (Figure 2).

### A validation experiment confirms the emergence of clonal bacterial strains that express population-level pulses of GFP in response to pulses of tetracycline

To check the robustness of the results reported by Ha et al.^34^, we conducted a follow-up experiment under different experimental conditions, in which deep-well plates were used instead of culture tubes (Methods). Overnight cultures of the pUC, CloDF13, pBR322, and p15A clones were plated for single colonies on LB+Kan+Tet agar, and 10 biological replicate populations were founded from individual colonies, for a total of 40 clones. This bottlenecking procedure was used to minimize the possibility that the observed pulses of GFP expression were due to competition between ancestral subpopulations of cells containing pure intracellular populations of *tetA*^*+*^ or *tetA*^−^ plasmids. The 10 clones for each plasmid treatment were split into two blocks of 5 clones per plasmid treatment and each block was passaged in deep-well plates by different experimenters (“G-block” and “R-block”). Three pulses of 10 μg/ml tetracycline were applied over the course of this experiment, and GFP fluorescence and OD600 were measured daily. Since 10 μg/ml tetracycline is lethal to ancestral clones that only contain one *tetA* copy in the chromosome, this experimental design further excludes the possibility that the observed pulses of GFP expression were due to competition between ancestral subpopulations of cells containing *tetA*^*+*^ or *tetA*^−^ plasmids.

The results of this experiment are shown in Figure 4F and Supplementary Figure 7. Importantly, the G-block and R-block used different air-permeable membranes to cover the deep-well plates used for culturing, resulting in significantly different means (Wilcox test: *p* < 10^−16^) and variances (Brown-Forsythe test: *p* < 10^−16^) in optical density for cultures grown in G-block versus R-block (Supplementary Figure 8). pUC clone 4 of G-block, CloDF13 clone 1 of R-block, and pUC clone 2 of R-block show large population-level pulses of GFP in response to pulses of tetracycline selection, while other clones show smaller GFP responses. Strikingly, limited dynamical changes are seen for many clones, especially those in R-block.

These results demonstrate that a further set of clones isolated from the clones used by Ha et al. shows biphasic behavior, in which some clones show a large GFP response to tetracycline selection, but others show a much weaker response. The biphasic behavior of clones isolated after further bottlenecking in this August 2025 experiment is consistent with two classes of clones: those containing plasmids saturated with the *tetA-gfp* transposon, and those that have maintained an intracellular balance of *tetA*^*+*^*/tetA*^−^ plasmids. Interestingly, the clones grown in the conditions of G-block show a stronger GFP response than those grown in the conditions of R-block. Therefore, the single clones used by Ha et al. produce daughter cells with wide variability in plasmid and *tetA-gfp* transposon copy numbers and frequencies; perhaps in part caused by fluctuations in plasmid copy number between growth in liquid media and solid agar for clone isolation, or differences in plasmid copy number dynamics depending on the different aeration conditions of the G-block and R-block of the August 2025 validation experiment.

## DISCUSSION

Here, we report the *de novo* evolution of clonal bacterial strains that modulate gene expression in response to pulses of tetracycline. Balancing selection stably maintains genetically diverse intracellular plasmid populations within single cells, due to the toxic effects of a *tetA* transposon. TetA toxicity creates a stable equilibrium between *tetA*^*+*^ and *tetA*^−^ plasmids, resulting in a biomolecular feedback control system. The intracellular balance of plasmids containing the *tetA-*transposon responds to antibiotic selection. The stability of the balancing selection regime and the magnitude of the GFP response increases with plasmid copy number. In the balancing selection regime, the distribution of plasmids within cells serves as an internal model of the environment, allowing the bacterial population to track environmental antibiotic concentrations.

The key property necessary for population-level feedback control is the maintenance of coexisting variants within a single cell by balancing selection. When this condition holds, the stable intracellular equilibrium among intracellular variants can be tuned by environmental selection. Such conditions can arise during the evolution of novel antibiotic resistances on multi-copy plasmids, in which the maintenance of ancestral and evolved TEM-1 β-lactamase alleles on multi-copy plasmids can overcomes evolutionary tradeoffs between ampicillin and ceftazidime resistance^35^.

The quasispecies framework that we use to describe the dynamics of intracellular *tetA*^*+*^/*tetA*^−^ plasmids can easily generalize to diverse intracellular populations of DNAs, RNAs, proteins, and organelles in other contexts. Our basic theoretical framework can apply to the evolution of novel antibiotic resistances on multicopy plasmids^35^, intracellular populations of extrachromosomal DNAs in highly aggressive cancers^36–39^, intracellular populations of circular RNA Obelisks^40,41^, mitochondrial heteroplasmy^42^, and heritable populations of proteins with diverse post-translational modifications, quaternary assemblies, or conformations^43,44^. Our mathematical model implies that the coexistence phase is suboptimal in constant environments, indicating that the emergence of intracellular quasispecies either occurs as an evolutionary response to transient selection pressures, or in response to persistent environmental fluctuations^35,45,46^. A quantitative understanding of how the variability in intracellular quasispecies may match rates of environmental fluctuations remains an open question, with implications for the fitness tradeoffs inherent in maintaining population diversity and phenotypic heterogeneity^47^. A key gap left by this work are the effects of simultaneous variation in the absolute numbers of plasmids per cell. The companion paper by Ha et al.^34^ addresses this question and its implications for quantitative microbiology.

Our results also have implications for systems and synthetic biology. Systems and synthetic biologists have largely drawn inspiration from electrical engineering and computer science in the construction of synthetic biological systems, so fundamental principles from evolutionary theory remain underappreciated. The implications of the Price equation for dynamically controlling bacterial adaptation have been overlooked by synthetic biologists constructing similar systems^18^. We address this by showing how the Price equation^48^ describes the behavior of adaptive systems constructed with intracellular plasmid populations^18^. Evolutionary processes are often omitted when engineering biomolecular circuits in living cells, and the rational engineering of autonomous evolutionary computing in living cells remains an overarching challenge for synthetic biologists^49^. Our findings demonstrate the value in bridging the gap between evolutionary and synthetic biology.

## Methods

### Strain construction protocol

Strain construction followed the protocols described in previous work^20,21^. All plasmids were transformed into strains using electroporation or chemical transformation using TSS buffer^50^. Following ref. ^20^, the helper plasmid pA004 was transformed into *E. coli* DH5α and *E. coli* K-12 MG1655. tetA-Tn5 mini-transposons were integrated into the host chromosome as follows. The mini-transposon plasmids contain the HK022 attP sequence. These plasmids were transformed into the host strain containing pA004 and inserted into the HK022 attB site on the *E. coli* chromosome by attB/P recombination.

### Nine-day evolution experiment with E. coli DH5α

The nine-day evolution experiment with *E. coli* DH5*α* was reported in previous work^21^; however, the gel and sequencing results reported here have not been published before.

#### Culture conditions

3mL cultures were grown in 16mL 17×100mm culture tubes at 37C in a 225-rpm shaking incubator at 37C. The cultures were propagated by 1:1000 daily serial dilution: 3μL of the Lysogeny Broth (LB) overnight cultures were used to inoculate 3mL LB + tetracycline and 3mL LB without tetracycline as a control.

#### Ancestral strains

The following *tetA-*Tn5 mini-transposons were integrated into *E. coli* DH5*α*. pB030 contains active Tn5 transposase outside of a mini-transposon containing *tetA* expressed under the strong J23104 promoter (Tn5+ *tetA++*). pB059 contains a mini-transposon containing *tetA* expressed under the strong J23104 promoter, but does not contain Tn5 transposase (Tn5− *tetA++*). pB020 contains active Tn5 transposase outside of a mini-transposon containing *tetA* expressed under the weak J23113 promoter (Tn5+ *tetA+*). The P15A plasmid pA031 was also transformed into these three strains. Altogether, six DH5*α* strains were evolved in LB with increasing tetracycline concentrations over time. These strains varied the presence of active transposase, the presence of an intracellular plasmid, and the strength of the promoter driving *tetA* expression.

#### Evolution experiment

3mL LB cultures were inoculated from glycerol stocks of the ancestral clones. The next day, 5× replicate populations were inoculated using 3μL of the overnight culture for each ancestral clone. Evolving populations were transferred nine times, increasing tetracycline concentrations in LB on each transfer (2, 4, 6, 8, 10, 20, 30, 40, 50 μg/mL). In total, 30 populations evolved in LB + tetracycline from the six ancestral strains. Another 30 populations were evolved in LB without tetracycline as a control. At the end of the experiment, 750 μL of each evolved population was mixed with 750 μL 50% glycerol in 2mL cryovials and stored at −80C.

#### Genomic analysis of transposon and plasmid copy numbers in endpoint clones from nine-day evolution experiment

Genomic DNA (gDNA) from endpoint mixed-population samples from the nine-day evolution experiment was extracted using the GenElute Bacterial Genomic DNA Kit (Sigma-Aldrich). gDNA was sent to SeqCenter (Pittsburgh, PA) for Illumina short-read genome sequencing. Variants were called using breseq version 0.3770 in polymorphism mode, using the following command-line flags: --polymorphism-minimum-variant-coverage-each-strand 4 -b 30 --maximum-read-mismatches 5. For the nine-day evolution experiment, a shell script called assemble-DH5a-genomes.sh was used to automate breseq runs and sequence data processing. Mini-transposon sequencing coverage data (for estimating copy-number change) was processed using the Python 3.12 script get-DH5a-expt-transposon-coverage.py. Analysis of transposon and plasmid copy numbers in the evolved population samples was conducted with an R 4.0 script called DH5a-expt-copy-number-analysis.R.

#### Detecting plasmid-based transposons by DNA gel electrophoresis

3mL LB + 10 μg/mL tetracycline + 50 μg/mL kanamycin cultures of single clones were inoculated from glycerol stocks and grown in in 16mL 17×100mm culture tubes at 37C in a 225-rpm shaking incubator at 37C. Plasmids were extracted from the culture and linearized with the restriction enzyme XhoI (New England Biolabs). Electrophoresis on a 1% agarose gel was used to separate and visualize plasmids with the transposon insertion (approximately 4.4 kilobases (kb)) from plasmids without the transposon (approximately 3 kb).

#### Measuring transposon ratio by real-time qPCR

Transposon and plasmid ratios were calculated by real-time qPCR following the protocol, reagents, and calibration described in ref. ^20^. Clones were cultured as described above, and 1 μl of a 1:10 culture dilution in PCR-grade H_2_O was used as the template for qPCR, which was performed using SuperReal PreMix probes (Thermo Fisher Scientific). The primers and probes used for qPCR are listed in Supplementary Tables 5 and 6 of ref. ^20^.

### Quasispecies model of transposon-plasmid evolutionary dynamics

We built a mathematical model to examine how plasmid copy number affects *tetA-GFP* copy number dynamics in the *tetA*-transposon system. We use the quasispecies model^29^, which has a physical interpretation as a model of continuous culture in an idealized turbidostat, such that the dilution rate is equal to the bulk growth rate (i.e. mean population fitness). The idea here is that the dilution rate increases as the population adapts, such that the total population density is conserved. Physically, this corresponds to a turbidostat in which the dilution rate instantaneously increases as the population increases in mean growth rate through evolutionary adaptation, such that the (optical) density of the culture stays constant.

The structural form of the quasispecies model is shown in Figure 2A, and a diagram of the quasispecies model for the *n* = 3 case is shown in Fig. 2B. The model involves *n+*1 subpopulations of bacteria: the first carries a copy of the *tetA* transposon on the chromosome and zero copies on the *n* plasmids in the cell; each additional subpopulation contains an additional copy of the *tetA* transposon on a plasmid, up to saturation where all *n* plasmids contain the *tetA* transposon. We are interested in the dynamics of these subpopulations due to selection (growth and dilution) and mutation (largely determined by plasmid segregation dynamics, with a small contribution due to transposition from the chromosome to plasmids).

See Supplementary Data 1 for an interactive Pluto computational notebook of the model written in Julia 1.11. This notebook can be run by installing and running Pluto.jl within Julia 1.11+ (see instructions at: https://plutojl.org/) and then opening the notebook using the Pluto web browser interface. Unless otherwise stated, the simulation results shown in Figures 3 and 4 use the following default parameter settings (arbitrary units): σ = 10.0 (this parameter determines the width of the fitness function) maximum growth rate *r*_*max*_ = 1.0,, Transposition Rate η = 0.001, Plasmid copy number *n* = 50, [Tet] concentration = 25.0.

#### Model assumptions

The population is modeled as a distribution over *tetA* copy numbers, ranging from 1 (found on the chromosome) to *n*+1, where *n* is the maximum plasmid copy number. We x_l_ therefore represent the population as a vector 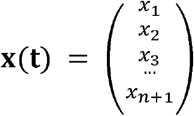. The sum of the *n*+1 entries of ***x****(t)* gives the total population size 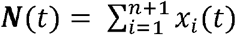.

#### Model assumptions I. growth dynamics

For a given tetracycline concentration, each subpopulation (*tetA* copy number class) has a growth rate, or *Malthusian fitness r*_*i*_. Let *g*_*i*_ be the log-fitness of *tetA* copy number class *I*, such that *g*_*i*_ = *log*(*r*_*i*_). We assume that there is some optimal *tetA* copy number given some tetracycline concentration. We assume that *r*_*i*_ is non-negative, with a single peak at the optimum *tetA* copy number for a given [*Tet*] concentration, so a natural choice is to define the following quadratic log-fitness function:

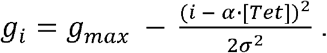

Here, *g*_*max*_ is the maximum log-growth-rate, *i* is the number of *tetA* copies in this strain, [*Tet*] is the tetracycline concentration, σ is a free parameter that determines the width of the quadratic function, and *α* is a unit-conversion factor with units of “*tetA* gene copies per unit [*Tet*]”. Without loss of generality we assume *α* = 1, so:

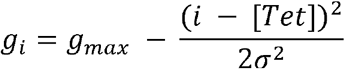

We assume fitness is non-negative, so we pass this *g*_*i*_ through an exponential function to get a Gaussian fitness function: *r*_*i = e*_^*gi*^. Then, the growth rates (Malthusian fitnesses) for each subpopulation is a vector 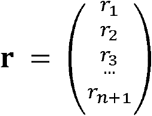. We then define the diagonal matrix 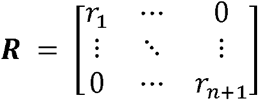 which represents how each subpopulation of the vector **x** grows based on **r**. Note that the average population growth rate (mean population fitness) of the whole population is 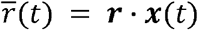

#### Model assumptions II. phenotype switching dynamics

Subpopulations switch phenotypes based on plasmid segregration during cell division. We assume that the total plasmid copy number is fixed per cell. However, the cells of each subpopulation can gain or lose plasmids containing the *tetA* transposon, based on sampling the plasmids of the parental cell with replacement. This assumption leads to a stochastic switching matrix based on binomial probabilities. Since the first index represents a state in which *none* of the plasmids have the transposon, we use a change-of-variable from zero-based indexing to one-based indexing, so that we can use the binomial formula (zero-based indexing) and then use the change-of-variable to use one-based indexing for the matrix. We define the entries of the stochastic transition matrix **S** with zero-based indexing:

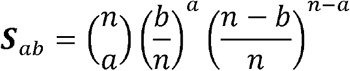

where *a* is the zero-based row-index (representing an offspring with *a* copies of the *tetA-*transposon) and *b* is the zero-based column-index (representing a parent with *b* copies of the transposon). Recall that *n* is the plasmid copy number, so 0 < *a* < *n* and 0 < *b* < *n*. We then use the following change of variables to define **S** with one-based indexing. The matrix is exactly the same— only the indexing the refer to entries is different:

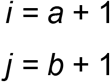

Here is an example of the 3-dimensional case:

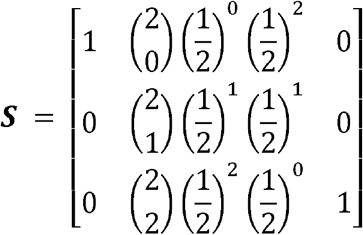

Mathematically, the subpopulation in which all plasmids have the transposon is an absorbing state of the Markov chain. However, the subpopulation in which *none* of the plasmids have the transposon is *not* an absorbing state, because transposons can jump from the chromosome to a plasmid. To model this, we assume a transposition rate *η*, and assume that *η* is small compared to the binomial transition probabilities so that it can be ignored in the other entries of the matrix (3-dimensional case shown):

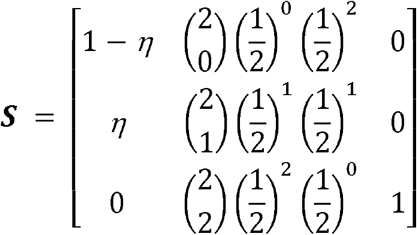

#### Model assumptions III. dilution dynamics

We assume that cells are diluted out at a rate equal to the bulk population growth rate, which is the average population growth rate (mean population fitness) of the whole population 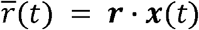.

#### Model assumptions IV. full dynamics

The growth and stochastic switching dynamics are combined into a matrix product **A = SR**, where **S** is the stochastic switching matrix and **R** is the diagonal growth-rate matrix. Including dilution, the full dynamics are modeled by the following matrix system of ODEs:

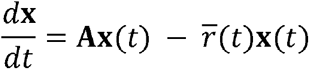

Note that 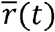 is a scalar and not a vector.

### Construction of barcoded E. coli K-12 MG1655 strains containing tetA-gfp mini-transposons

A *tetA-gfp* Tn5 mini-transposon vector constructed in previous work^20^ named pB31 was modified to contain random barcodes outside of the Tn5 IE and OE end sequences, following the protocol described in ref. ^20^. The barcoded transposon vector library is called B31-N20, where N20 refers to two 10 bp randomized sequences (the random barcode) flanking the AscI restriction enzyme cut site on the B31-N20 transposon vector. Briefly, the transposon vector backbone was generated by linearizing the *tetA-gfp* transposon vector (pB31) using PCR amplification. A DNA fragment containing the random barcode was synthesized by Integrated DNA Technologies. The randomized barcode DNA fragment was assembled into the transposon vector backbone by Gibson Assembly cloning. The library of barcoded transposon vectors was generated by transforming the Gibson Assembly reaction into electrocompetent cells, allowing an overnight outgrowth of the transformed cells, and then collecting the barcoded transposon vector library by plasmid miniprep the following morning. Transposon vectors were then integrated into the chromosome of *E. coli* K-12 as described above, to yield a collection of isogenic strains containing chromosomal *tetA-gfp* Tn5 mini-transposons, each containing unique barcodes in the chromosome and outside the *tetA-gfp* mini-Tn5 transposon itself. This process was repeated to generate isogenic barcoded K-12 strains containing the pUC plasmid pA18 and the p15A plasmid pA31. Sequence barcodes were verified by Sanger sequencing before further experiments.

### Isolation of barcoded tetA-gfp transposon clones for pulsed tetracycline experiments

3mL LB + 35 μg/mL chloramphenicol + 50 μg/mL kanamycin cultures of the following ancestral barcoded K-12 clones with *tetA-gfp* transposons integrated in the chromosome (but not present on plasmids) were inoculated from glycerol stocks and grown overnight in 16mL 17×100mm culture tubes at 37C in a 225-rpm shaking incubator at 37C: RM7.107.38 (containing pUC plasmid), RM8.24.2 (containing CloDF13 plasmid), RM8.27.4 (containing pBR322), RM7.107.3 (containing p15A), RM8.31.1 (containing pSC101). The next day, 3 μL of overnight culture was transferred to fresh 3mL LB + 2μg/mL tetracycline cultures and grown overnight. The following day, 3 μL of overnight culture was transferred to fresh 3mL LB + 10μg/mL tetracycline cultures and grown overnight. The next day, cultures were dilution plated for single colonies on LB + 10 μg/mL tetracycline agar plates and grown in a static incubator at 37C. The next day, single colonies were picked and grown in 3mL LB + 10 μg/mL tetracycline cultures. 750 μL of each evolved clone was mixed with 750 μL 50% glycerol in 2mL cryovials and stored at −80C. The following evolved clones were isolated by this procedure and used for pulsed tetracycline experiments: RM8.35.1 (pUC clone), RM8.35.2 (CloDF13 clone), RM8.35.3 (pBR322 clone), RM8.35.4 (p15A clone), RM8.35.5 (pSC101 clone). Specifically, the main pulsed tetracycline evolution experiments carried out by Ha et al.^34^ (2025) (and the follow-up August 2025 experiment described in this work) used the RM8.35.1, RM8.35.2, and RM8.35.3 strains. The August 2025 experiment also used the RM8.35.4, although gel electrophoresis experiments indicate that this clone contained an empty p15A plasmid lacking the *tetA-gfp* transposon.

### Comparing ancestral (empty) and evolved (containing tetA-gfp transposons) plasmids by DNA gel electrophoresis

3mL LB + 35 μg/mL chloramphenicol + 50 μg/mL kanamycin cultures of the following ancestral K-12 *tetA-gfp* transposon clones were inoculated from glycerol stocks and grown overnight in 16mL 17×100mm culture tubes at 37C in a 225-rpm shaking incubator: RM7.107.38 (pUC ancestral strain), RM8.24.2 (CloDF13 ancestral strain), RM8.27.4 (pBR322 ancestral strain), RM7.107.3 (p15A ancestral strain), RM8.31.1 (pSC101 ancestral strain). 3mL LB + 10 μg/mL tetracycline cultures of the following evolved K-12 *tetA-gfp* transposon clones were inoculated from glycerol stocks and grown overnight in the same conditions: RM8.35.1 (pUC clone), RM8.35.2 (CloDF13 clone), RM8.35.3 (pBR322 clone), RM8.35.4 (p15A clone), RM8.35.5 (pSC101 clone). Plasmids were extracted from cell pellets of 1mL of overnight culture and linearized with the restriction enzyme XhoI (New England Biolabs). Electrophoresis on a 1% agarose gel was used to separate and visualize plasmids with the transposon insertion (approximately 4.4 kilobases (kb)) from plasmids without the transposon (approximately 3 kb). Gel electrophoresis indicated that the *tetA-gfp* transposon mobilized onto the pUC, CloDF13, and pBR322 plasmids in RM8.35.1-3, but did not mobilize onto the p15A plasmid in RM8.35.4 (Supplementary Figure S5).

### August 2025 pulsed tetracycline evolution experiment

3mL LB + 10 μg/mL tetracycline + 50 μg/mL kanamycin cultures of RM8.35.1-4 (containing pUC, CloDF13, pBR322, and p15A plasmids, respectively) were grown in standard conditions (37C in a 225-rpm shaking incubator) overnight. The next day, cultures were dilution plated for single colonies on LB + 10 μg/mL tetracycline + 50 μg/mL kanamycin agar plates and grown in a static incubator at 37C. The next day, 10 colonies from each agar plate were picked and used to inoculate 1mL LB + 10 μg/mL tetracycline + 50 μg/mL kanamycin cultures in two deep-well plates (5 independent colonies per plasmid treatment per plate) in a shaking plate incubator at 37C and 700 rpm. The next day (Day 0 of this experiment), 1 μL of each culture on each plate was transferred into fresh 1mL LB + 10 μg/mL tetracycline wells (Kan− treatment) and a paired set of 1mL LB + 10 μg/mL tetracycline + 50 μg/mL kanamycin wells (Kan+ treatment). The paired sets of cultures were propagated in 1 mL LB and 1 mL LB+Kan for 12 transfers, where pulses of 10 μg/mL tetracycline were added on Days 0, 3, and 10. Cultures were transferred as 1:1000 dilutions (1μL of overnight culture to fresh media) for the first three transfers (days 0, 1, 2, 3), and as 1:10,000 dilutions (10μL of overnight culture into an intermediate deep-well dilution plate containing 1mL LB, and then 10μL of this dilution into the fresh deep-well plate for the experiment) for the rest of the experiment. Every day, OD600 and GFP fluorescence was measured on 100uL of culture in a Corning Costar black clear-bottom plate in a Tecan Infinite 200 Pro plate reader. Note that the two deep-well plates represented two blocks of the experiment (G-block and R-block) and used different air-permeable membranes to seal the plate: Sigma-Aldrich Breathe-Easy sealing membrane #Z380059 for G-block and Millipore Sigma AeraSeal film #A9224-50EA for R-block.

## Acknowledgements

We thank Yi Yao for constructing the barcoded *tetA-gfp* transposon system. We thank Kyeri Kim, Zhengqing Zhou, and the members of the You lab for valuable comments and discussions. We thank Bin Li and the Duke Cancer Institute Flow Cytometry core for technical assistance. We thank Richard Lenski and Jeffrey Barrick for feedback on an earlier draft. This work was partially funded by the National Institutes of Health (L.Y., R01AI125604, R01GM098642, and R01EB031869) and the New Jersey Agricultural Experiment Station (startup funds to R.M.).

## SUPPLEMENTARY INFORMATION

**Supplementary Data 1: Pluto computational notebook, written in the Julia programming language, allowing for user interaction with the plasmid quasispecies model**.

**Supplementary Figure S1.**
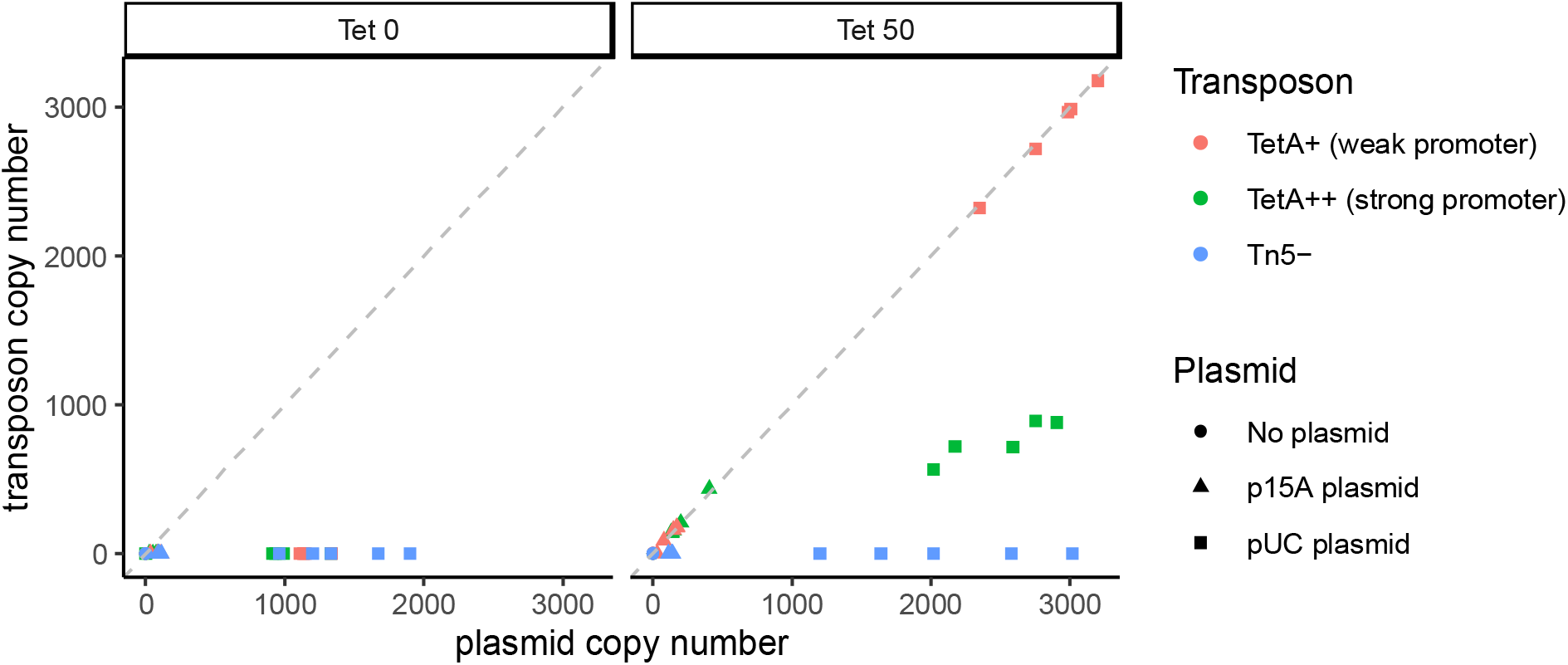
TetA++ transposons do not saturate pUC plasmids in a 9-day evolution experiment.

**Supplementary Figure S2.**
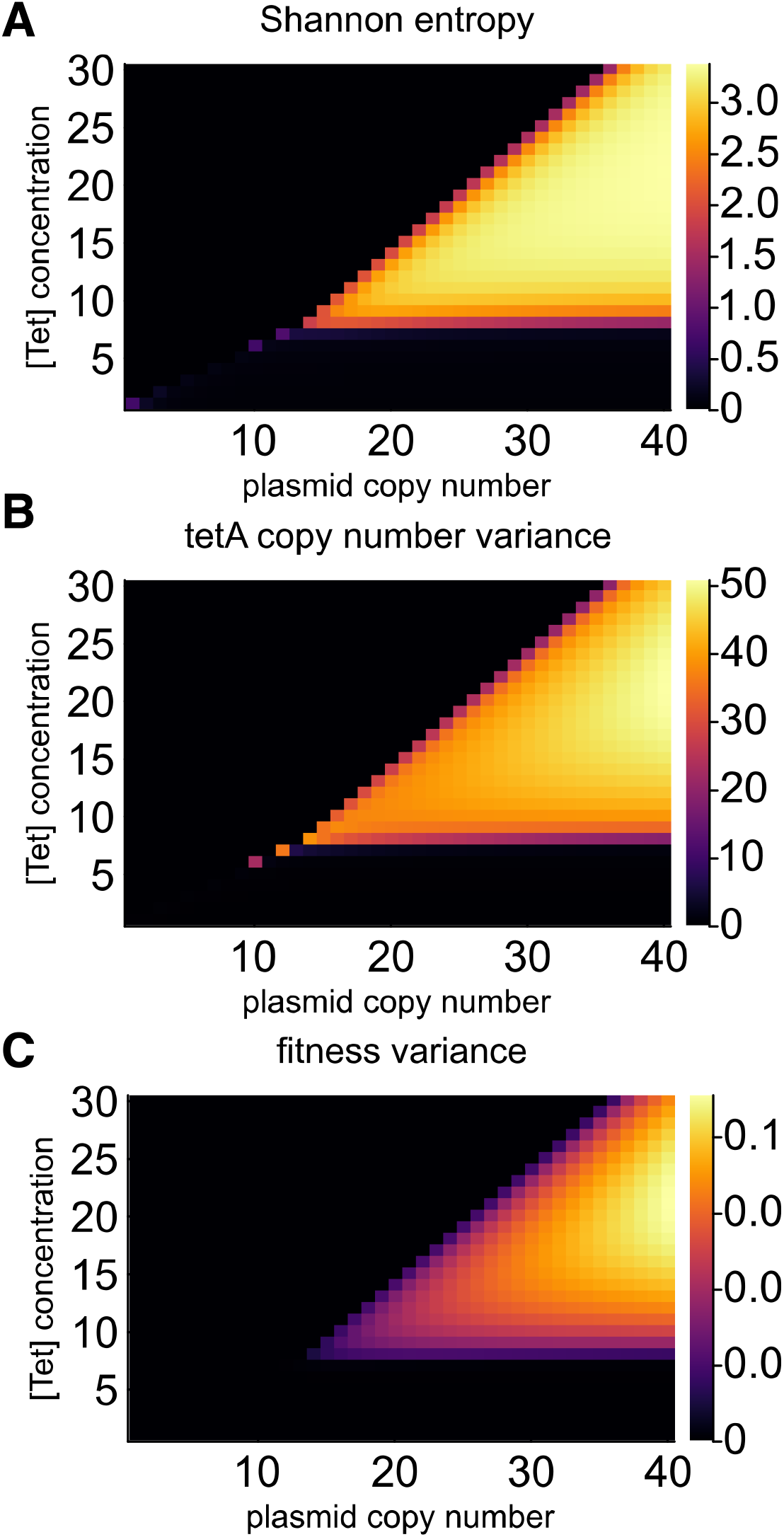
Phase diagram reveals the emergence of intracellular plasmid coexistence. The sparse dots on the diagonal outside of the coexistence region correspond to the unstable coexistence of subpopulations of cells containing only monomorphic populations of *tetA*^*+*^ or *tetA*^−^ plasmids due to comparable fitnesses (resulting in positive Shannon entropy and *tetA* copy number variance, but near-zero fitness variance). A) Shannon entropy in the phase diagram. B) *tetA* copy number variance in the phase diagram. C) Fitness variance in the phase diagram.

**Supplementary Figure S3.**
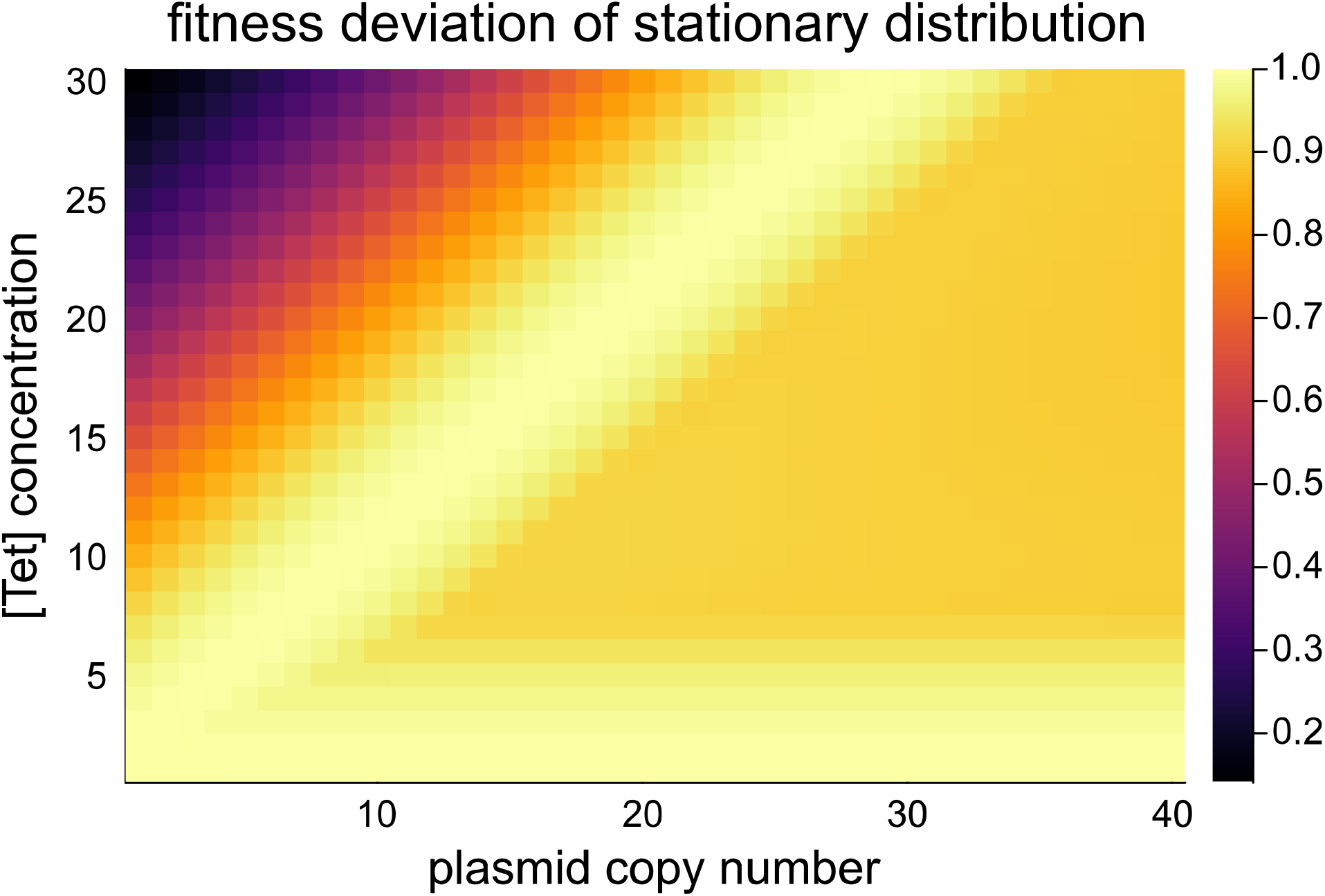
Coexistence phase has suboptimal fitness in a constant environment.

**Supplementary Figure S4.**
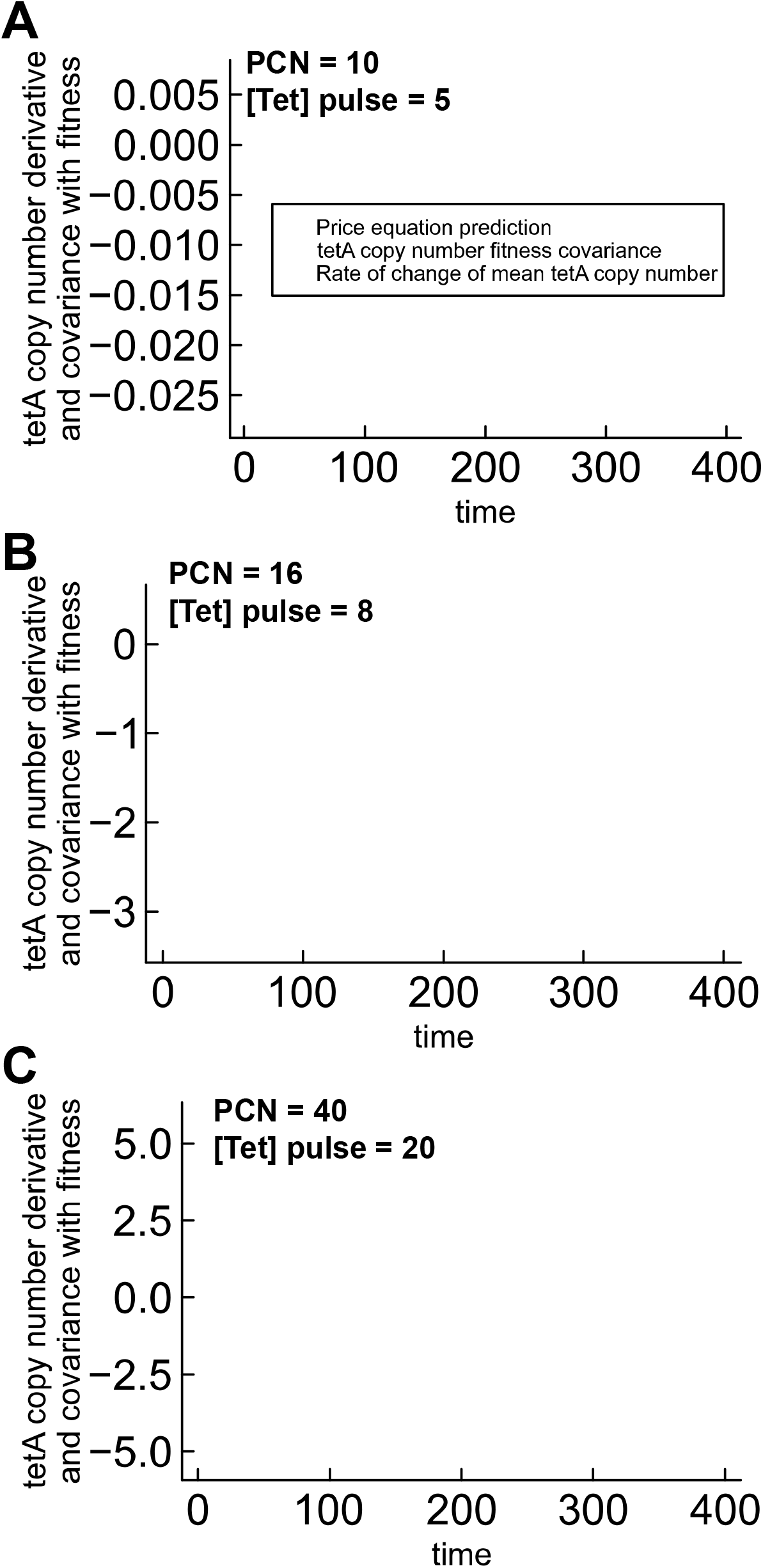
Price equation predicts quasispecies dynamics in the cases shown in the phase diagram.

**Supplementary Figure S5.**
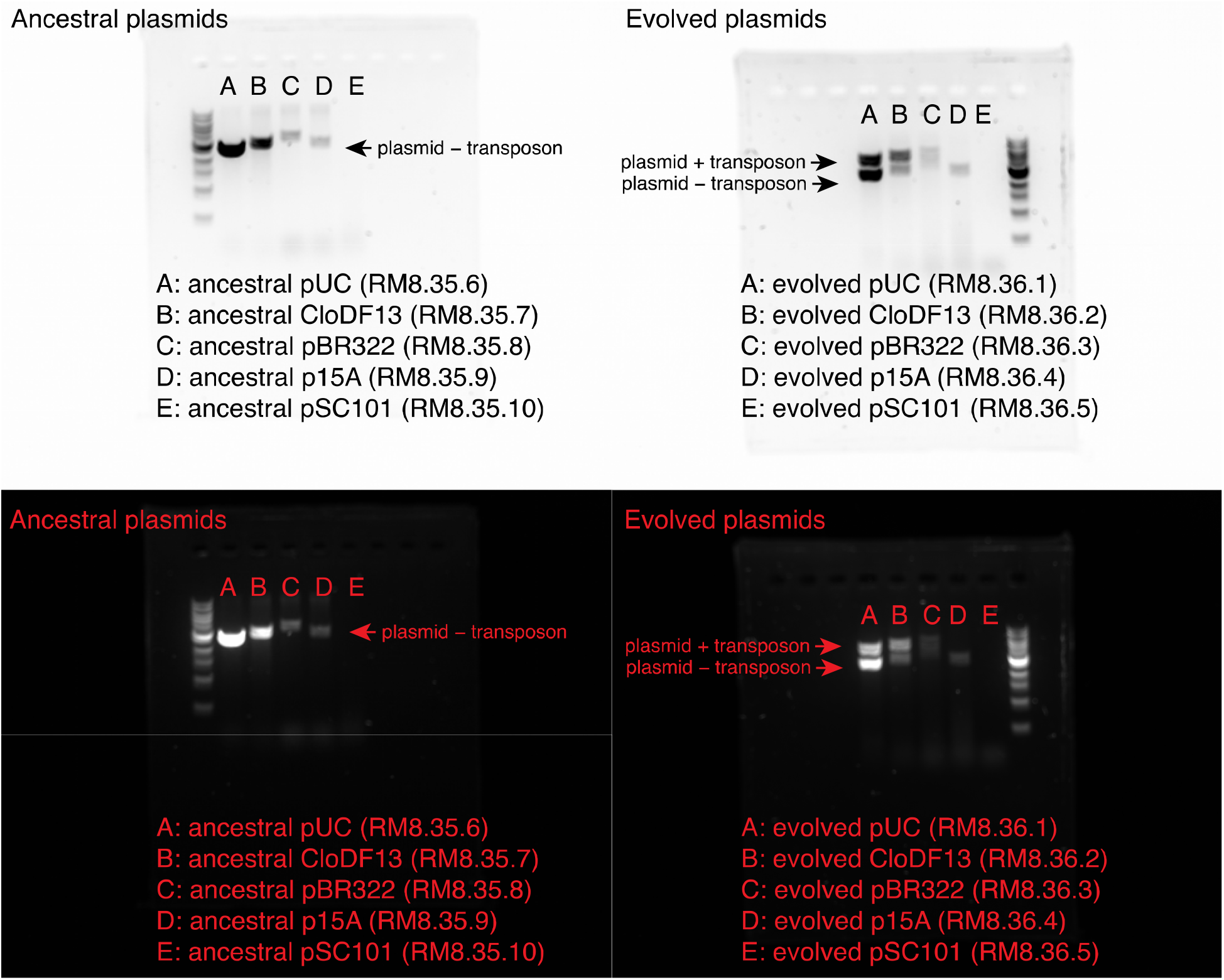
DNA electrophoresis shows coexistence of *tetA*^+^ and *tetA*^−^ plasmids in evolved clones containing pUC and CloDF13. The top and bottom panels show the same gel images, but with inverted color channels.

**Supplementary Figure S6.**
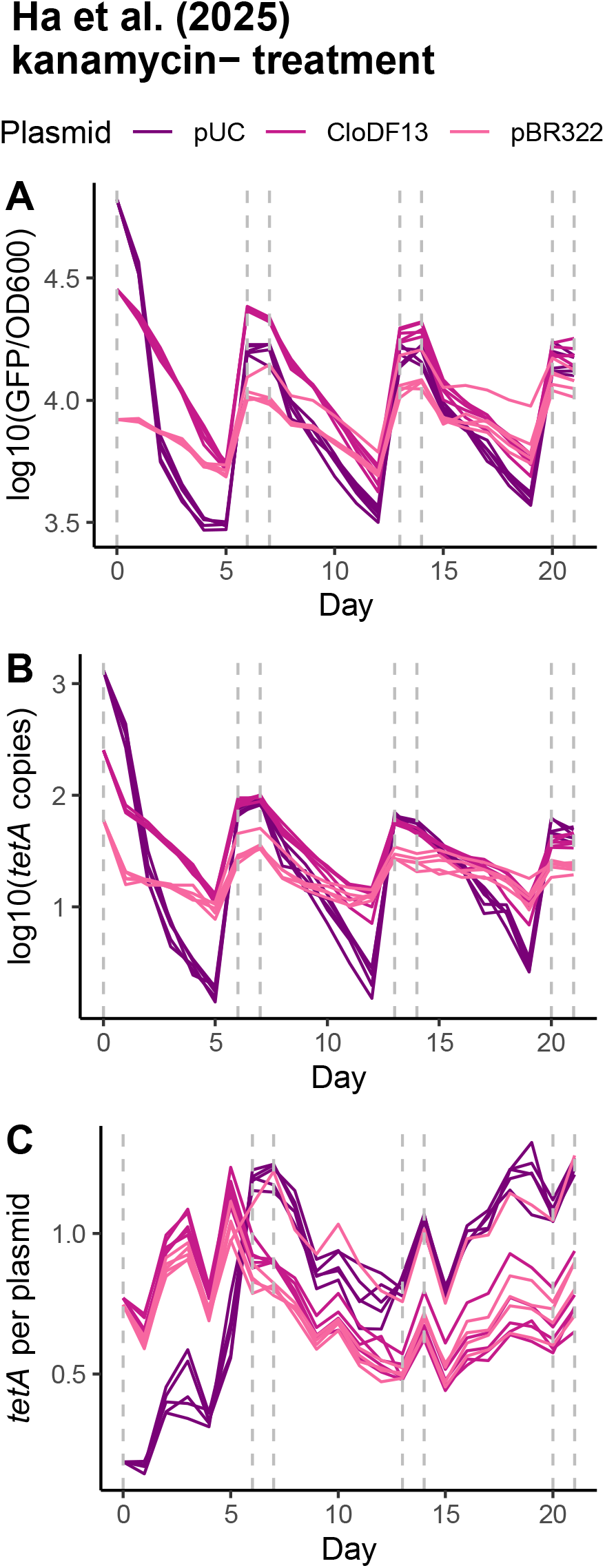
Experiments carried out by Ha et al. (2025) show that intracellular *tetA*^+^ plasmid copy number responds to tetracycline for clones containing pUC, CloDF13, and ColE1, in the kanamycin− treatment. Dashed gray lines indicate timing of tetracycline pulses.

**Supplementary Figure S7.**
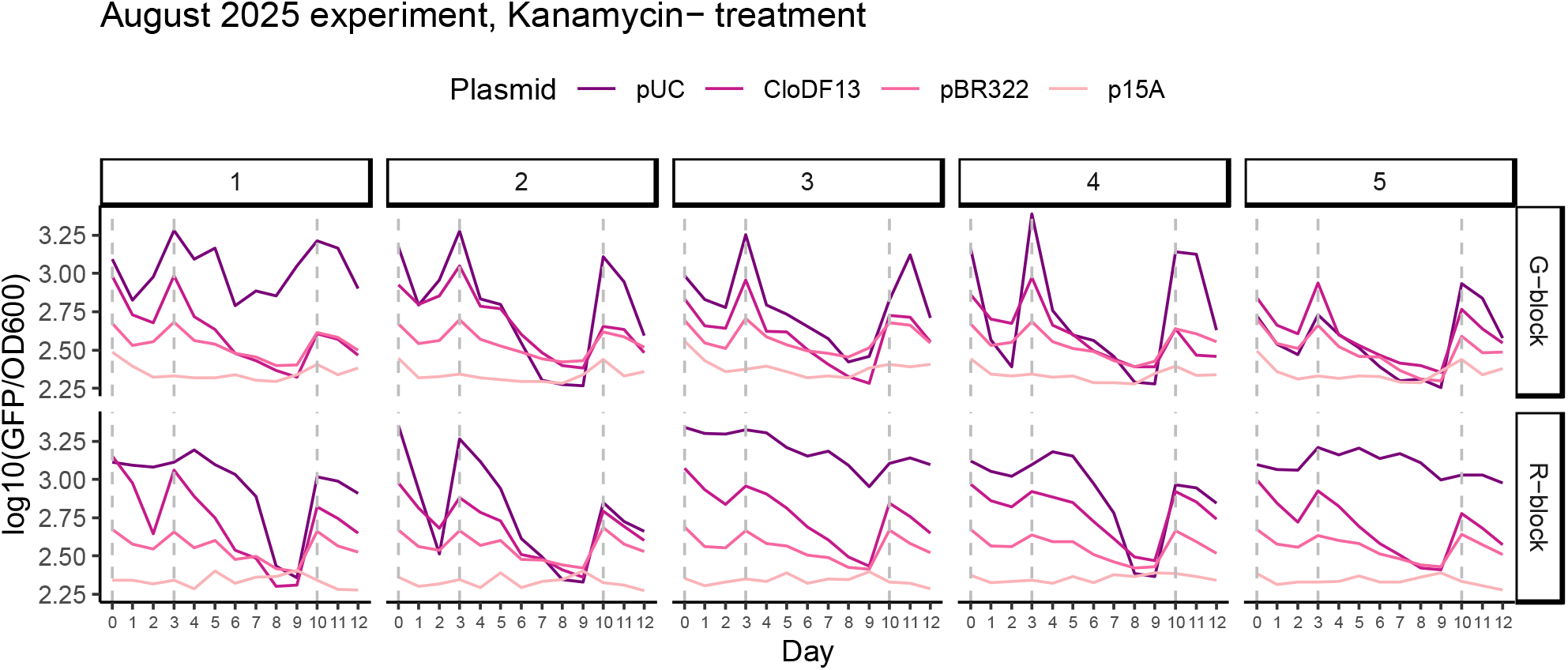
Follow-up validation experiment on clones isolated through a second bottleneck shows that *tetA-gfp*^+^ plasmid copy number responds to tetracycline in the kanamycin− treatment. Dashed gray lines indicate timing of tetracycline pulses.

**Supplementary Figure S8.**
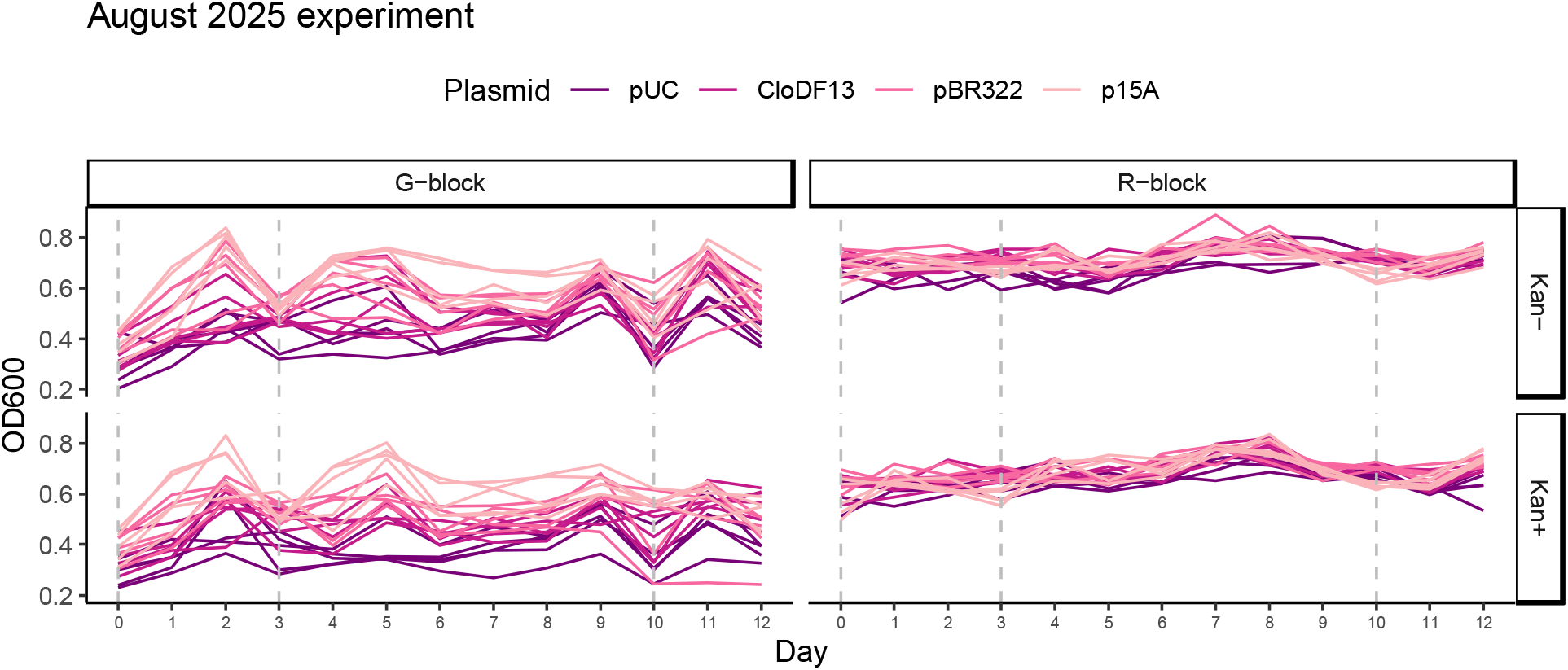
Choice of air-permeable membrane between the G-block and R-block of the August 2025 validation experiment results in a significant difference in mean and variance in OD600 over the course of the experiment. Dashed gray lines indicate timing of tetracycline pulses.

